# Computational investigation of *cis*-1,4-polyisoprene binding to the latex-clearing protein LcpK30

**DOI:** 10.1101/2023.06.26.546638

**Authors:** Aziana Abu Hassan, Marko Hanževački, Anca Pordea

## Abstract

Latex clearing proteins (Lcps) catalyze the oxidative cleavage of the C=C bonds in *cis*-1,4-polyisoprene (natural rubber), producing oligomeric compounds that can be repurposed to other materials. The active catalytic site of Lcps is buried inside the protein structure, thus raising the question of how the large hydrophobic rubber chains can access the catalytic center. To improve our understanding of hydrophobic polymeric substrate binding to Lcps and subsequent catalysis, we investigated the interaction of a substrate model containing ten carbon-carbon double bonds with the structurally characterized LcpK30, using multiple computational tools. Prediction of the putative tunnels and cavities in the LcpK30 structure, using CAVER-Pymol plugin 3.0.3, fpocket and Molecular Dynamic (MD) simulations provided valuable insights on how substrate enters from the surface to the buried active site. Two dominant tunnels were discovered that provided feasible routes for substrate binding, and the presence of two hydrophobic pockets was predicted near the heme cofactor. The larger of these pockets is likely to accommodate the substrate and to determine the size distribution of the oligomers. Protein-ligand docking was carried out using GOLD software to predict the conformations and interactions of the substrate within the protein active site. Deeper insight into the protein-substrate interactions, including close-contacts, binding energies and potential cleavage sites in the *cis*-1,4-polyisoprene, were obtained from MD simulations. Our findings provide further justification that the protein-substrate complexation in LcpK30 is mainly driven by the hydrophobic interactions accompanied by mutual conformational changes of both molecules. Two potential binding modes were identified, with the substrate in either extended or folded conformations. Whilst binding in the extended conformation was most favourable, the folded conformation suggested a preference for cleavage of a central double bond, leading to a preference for oligomers with 5 to 6 C=C bonds, as shown by experimental data. The results provide insight into further enzyme engineering studies to improve catalytic activity and diversify the substrate and product scope of Lcps.

**Author summary:** Rubber materials are very important in our everyday life, but also lead to a high amount of rubber waste for which there is no sustainable solution. The enzymatic degradation of diene rubbers is an attractive option to revalorise these materials once they reach end of life. Latex clearing proteins (Lcps) have been shown to degrade polyisoprene rubber, however rates of degradation are low and the product is a mixture of oligomers. Enzyme engineering is necessary to develop a useful process leading to valuable materials, but it requires a thorough understanding of how substrate and protein interact. This is difficult to achieve when the substrates are large molecules with conformational flexibility. Here, we employed multiple computational tools to understand the interaction between LcpK30 and its flexible polyisoprene substrate. Our results show that the substrate can access the active site through two hydrophobic tunnels, which can also serve as product exit pathways. The substrate binds in a hydrophobic pocket near the heme, which determines the size of the oligomeric products. We identified two potential binding modes for the substrate and characterized the hydrophobic contacts responsible for protein-substrate complexation in LcpK30. These results shed light on future enzyme engineering investigations to enhance catalytic activity and broaden the substrate and product range of Lcps.

## Introduction

Rubber materials are essential in everyday life, with more than 25 million metric tons of natural and synthetic rubber being produced and consumed yearly [1]. This results in a high amount of rubber waste, which is challenging to recycle or decompose, thus creating a serious environmental concern once the material reaches the end of life. An economically and environmentally sustainable strategy for rubber waste disposal is yet to be established. The degradation of rubber polymers mainly involves oxidative cleavage of C=C bonds in the polymer backbone and has been reported with organic or organometallic catalysts, which are limited by the use of rare metals (e.g. Grubbs catalyst) and harsh reagents [2]. An attractive alternative is enzymatic rubber degradation using oxidative enzymes. Rubber oxygenases (RoxA and RoxB), isolated from Gram-negative bacteria and latex clearing proteins (Lcps), isolated mostly from Gram-positive bacteria, have been shown to degrade both natural and synthetic *cis*-1,4-polyisoprene, with the latter enzymes being more extensively studied [3]. The products are functionalized oligoisoprenoids carrying a carbonyl group (ketone and aldehyde) at each end (Fig 1a). This cleavage preserves the polyisoprene structure and gives access to lower molecular weight products, which can be either further metabolized or used as materials with potentially interesting properties [4]. Thus, rubber degrading enzymes offer exciting perspectives for polymer degradation within a circular economy and have been extensively studied in the past few years.

**Fig 1.**
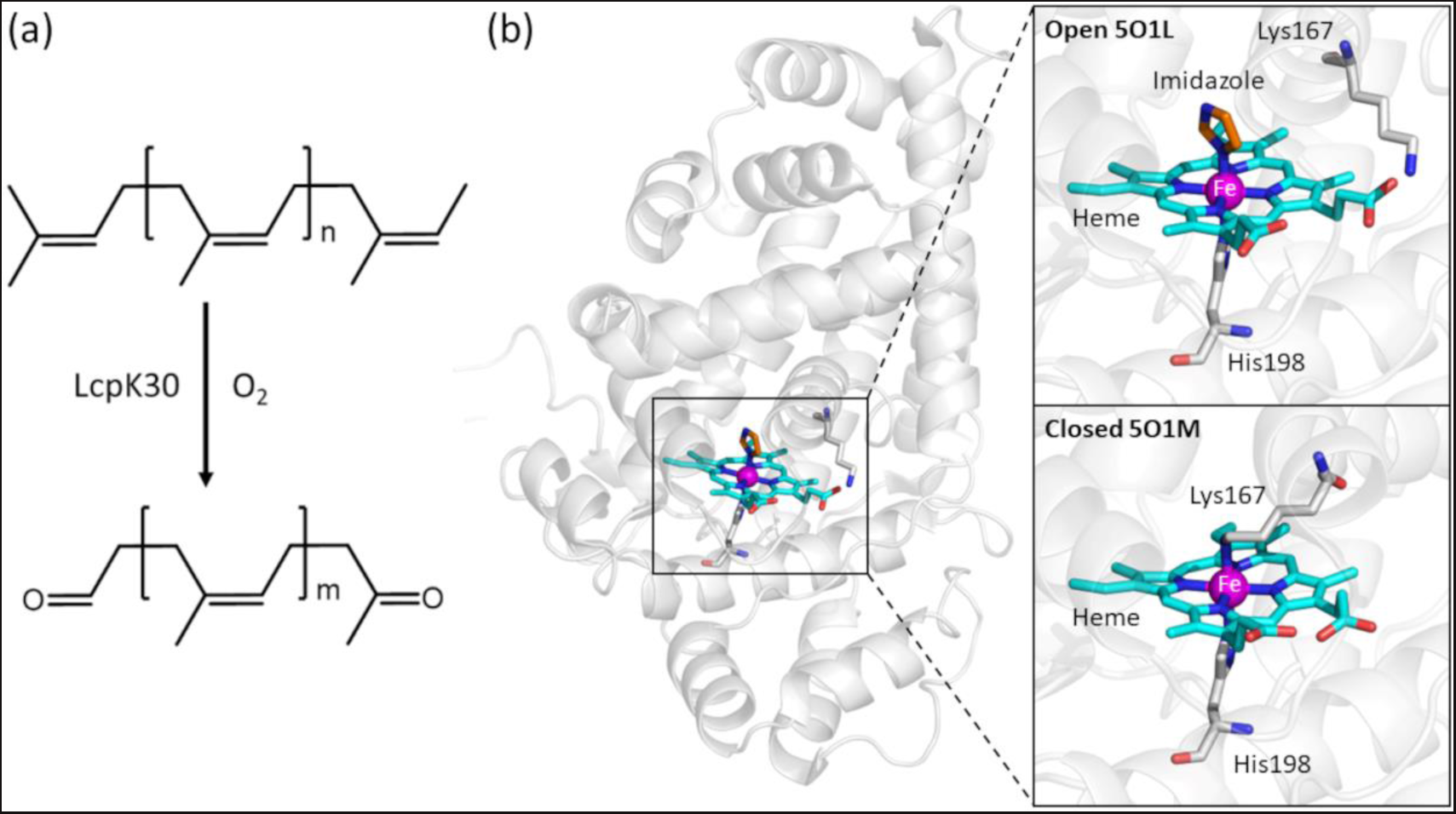
Cleavage of *cis*-1,4-polyisoprene catalyzed by LcpK30 and its crystal structure. (a) Oxidative cleavage of *cis*-1,4-polyisoprene catalyzed by LcpK30. (b) Crystal structure of LcpK30 (PDB ID: 5O1L) with a close view of heme in the active site. Coordination of an imidazole molecule to the distal site of heme fixes the protein structure in its open conformation (PDB: 5O1L), while the coordination by the sidechain of Lys167 forces the closed state of the enzyme (PDB: 5O1M).

A range of Lcps have been identified from several organisms [5–9] and were shown to be *b*-type cytochromes with a non-covalently bound heme cofactor. The crystal structure of LcpK30 has been determined to possess a globin-like fold, showing the heme to be coordinated to His198 (Fig 1b) [10]. The oxidative cleavage of polyisoprene by Lcp occurs in an endo-type mechanism and results in a mixture of oligoisoprenoids with different lengths, from C20 to C65 (3 to 12 isoprene units) [3,11–13]. A defined distribution of degradation products is observed, with a preference for 5-6 (for Lcp1VH2) or 5-8 (for LcpK30) isoprene repetitive units and a decrease in the concentrations for smaller and larger products [14,15]. Quantum mechanics/molecular mechanics (QM/MM) calculations suggested that the addition of the heme-bound dioxygen to the C=C bond triggers its cleavage via a dioxetane intermediate [16]. This mechanism is similar to two other heme-based dioxygenases, indoleamine and tryptophan 2,3-dioxygenases.

Structural and spectroscopic analysis of LcpK30 suggested the existence of two conformational states, closed and open states [10]. In the open state, a continuous hydrophobic substrate channel passes next to the heme group and allows the substrate to access the active site, whilst, in the closed state, the channel is blocked by coordination of the heme to Lys167. This difference is most visible in the conformation of the active site helix (also known as helix E),[10] comprised of residues 160-176. The helix is more structured in the closed state compared to the open state, in which this helix is split into two fragments (see S1 Fig in the Supporting Information (SI) for details). Most enzymes that react with polymeric substrates, such as PETases (with PET substrates) and lytic polysaccharide monooxygenases, LPMOs (with polysaccharide substrates) contain exposed active sites, which allow easy contact with the polymer chains of the substrate [17,18]. Intriguingly, the catalytic heme cofactor in Lcps is buried within a hydrophobic pocket, which raises the question of how the insoluble hydrophobic rubber chains can access the active site.

Despite an increased understanding of the structure and catalytic mechanism of Lcps, a useful process for enzymatic rubber degradation has not been achieved. The enzymatic degradation rates remain low, and the range of oligoisoprenoid products is too heterogeneous to be convenient for applications as materials. Furthermore, the enzymatic degradation is limited to *cis*-1,4-polyisoprene rubber, whilst the degradation of other diene rubbers remains largely underexplored. These challenges could potentially be addressed through protein engineering approaches, which require an understanding of the interaction between substrate and protein.

In this study, we used a computational modelling approach starting from the LcpK30 crystal structure in open conformation to understand the interaction between LcpK30 and its flexible polyisoprene substrate. Firstly, the potential access pathways of *cis*-1,4-polyisoprene to the active site of LcpK30 were explored using the CAVER PyMol plugin 3.0.3 and fpocket, to identify relevant hydrophobic tunnels and pockets that are large enough to accommodate the substrate [19]. We then carried out supramolecular docking of a C50 substrate model containing 10 repeating isoprene units (C_50_H_82_), which was deemed sufficiently large to provide valuable insights into the location, conformations and interaction of the polymer chains with the enzyme. Given the challenge to obtain precise information from the docking of long alk(en)yl chains without specific pharmacophores, we used molecular dynamics (MD) simulations to provide a better understanding of the stability and the nature of the predicted substrate poses and conformational changes of the protein.

## Results and discussion

### Identification of tunnels and pockets in LcpK30

Putative tunnels were identified within the crystal structure of LcpK30 in the open state, using the CAVER-PyMOL plugin to obtain a general overview of potentially important pathways for the substrate access from the surface to the active site. This plugin provides a graphical interface for setting up the calculation and allows interactive visualization of tunnels or channels in protein static structures [20]. A total of four tunnels were identified, and their lengths, radii, and bottlenecks (narrowest parts of the tunnels) were determined (see S1 Table and S2 Fig in the SI for details on the tunnel features). Tunnels 1 and 2 had the highest throughput and, thus, the highest probability to be used as routes for the transport of substrates (S1 Table and Fig 2a). Near the heme cofactor in the active site, these tunnels were lined up by residues known to be of high importance for LcpK30 activity: Glu148, Arg164, Lys167 and Thr168.[10,12] Other, more surface-exposed residues interacting with the tunnels included Trp82, Thr83, Arg84, Ala151, Val152, Gly157, Gly158, Ala159, Asp163, Ile165, Ala166, Ala169, Arg170, Leu171, Asp174, His184, Gly185, Ser186, Val189, Thr190, Lys193, Thr194, Val197, His198, Thr230, Glu392, Gly393, Arg394, Arg395, Ile396, Ala397, Ile398, Asp399, and Pro401. The radius of putative tunnels ranged between approximately 1 Å and 2.8 Å, with the bottleneck (0.96 Å) located near the heme and the wider radius found towards the outer surface of the protein.

**Fig 2.**
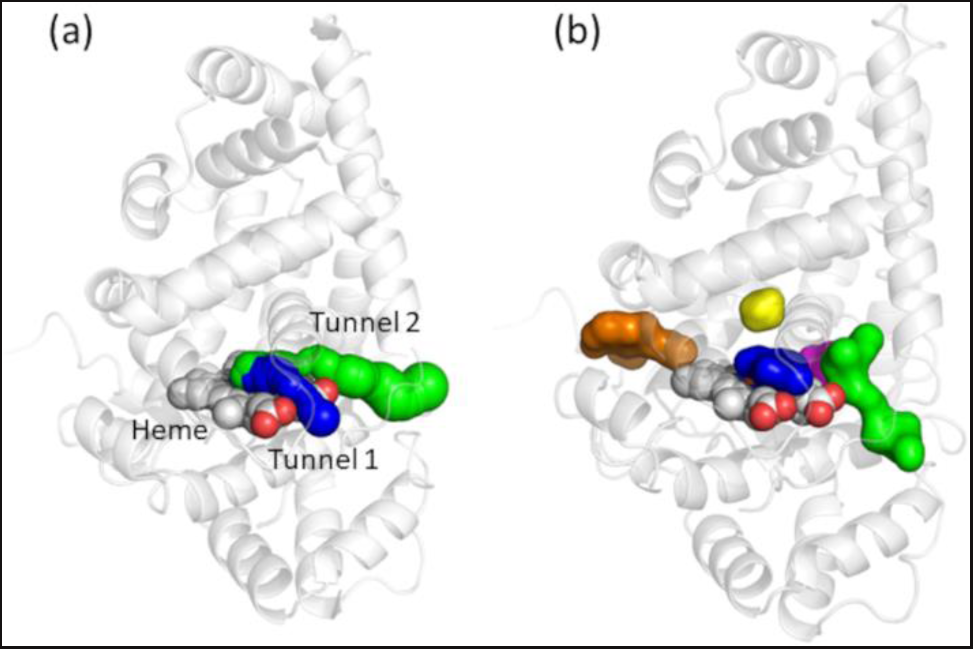
Tunnel mapping in LcpK30. (a) Two dominant tunnels identified by CAVER-Pymol plugin 3.0.3 (tunnel 1 in blue and tunnel 2 in green); (b) five detected pockets or cavities closest to heme in the LcpK30 static X-ray structure identified by automated geometry-based fpocket tool. The pockets are pocket 1 (green), pocket 3 (blue), pocket 4 (orange), pocket 9 (magenta) and pocket 12 (yellow).

The average radii throughout both tunnels were smaller than the estimated width of the *cis*-1,4-polyisoprene substrate molecule (see S2c Fig in SI), which suggested that significant conformational changes of the protein are required to accommodate the hydrophobic chains of the large polymeric substrates through this gateway. Alternatively, these tunnels are also suited for transporting smaller oxygen to the heme. We observed a significant difference in the length of the tunnels from the starting point to the protein surface, with tunnel 2 being about twice longer compared to tunnel 1. Whilst a shorter tunnel such as tunnel 1 may generally give quicker access of substrates to the active site, in this case, where the substrate is a long polymer, the longer tunnel 2 may provide more opportunity for interaction with a polymeric substrate. Furthermore, tunnel 2 has a wider opening (2.8 Å) at the protein surface (Glu392, Gly393, Arg394, Arg395, Ile396) which agrees well with the estimated thickness of *cis*-1,4-polyisoprene and which could represent a potential gateway for the substrate access to the active site.

We also predicted the important pockets (protein cavities) in the active site of the enzyme that could accommodate and position a hydrophobic substrate near the heme. Using an automated geometry-based fpocket tool that decomposes a 3D protein into Voronoi polyhedrals, we extracted the pockets from the protein surface using alpha spheres (see Methods for details). A total of 24 pockets were detected throughout LcpK30. However, only five pockets (pocket 1, 3, 4, 9 and 12) were considered relevant for substrate binding due to their near proximity to heme and the active site (Fig 2b). Out of these five, two pockets were especially interesting since they are located at a distal site directly above the heme cofactor: a larger pocket 3, with a pocket volume of 173.52 Å^3^ and a smaller pocket 12, with a volume of 77.21 Å^3^. These two pockets also had a relatively high hydrophobicity score, indicating that they could accommodate the lipophilic *cis*-1,4-polyisoprene ligand in its bound state. Interestingly, these cavities were also occupied by imidazole molecules in the crystal structure. The residues surrounding these two pockets are Ser138, Ser142, Glu148, Val152, Arg164, Lys167, Thr168, Leu171, Thr230, Ser233 and Leu234, and include the same residues lining up the tunnels identified earlier using CAVER- PyMOL plugin. Taken together, the information about the tunnels and pockets calculated on the X-ray structure of LcpK30, provided a better picture on possible entry pathways that lead from the surface into the hydrophobic active site of the protein interior and gave an indication of the space where ligand might bind.

Since the intrinsically dynamic nature of the protein could largely influence the pathways and cavities through the protein, we carried out extensive MD simulations of LcpK30 to obtain more information, specifically on the conformational changes that could influence binding of the substrate to LcpK30. Using our bonded model of heme, we were able to simulate the catalytically active state of the histidine-coordinated cofactor with O_2_ bound to the iron metal centre during relatively long simulation timescales. The calculated root-mean-square deviation (RMSD) of the C-alpha backbone atoms from the reference X-ray structure remained stable (1-1.5 Å) for 500 ns in all performed simulations (S3a Fig). The RMS fluctuation analysis of the C-alpha atoms revealed conformational changes of the protein linked to residues 155-160 from the loop near heme and residues 392-396 at the C-terminus (S3b Fig). Similar fluctuations were also found in the enzyme from the experimental crystallographic B-factor (S4 Fig). We further characterized these conformational changes as the loop rotation containing Arg394, which occasionally replaces Lys167, forming the salt bridge with the carboxylate group of heme during the simulation. Taking into account these dynamic features of the protein, we hypothesize that the structural adjustment of LcpK30 could facilitate the entry of substrate and exit of oligoisoprenoid products through the gateway comprised of C-terminus loop 392-396 and flexible loop 155-160.

We calculated and analyzed the access tunnels in dynamic LcpK30 structures obtained from MD simulations, using the CAVER-Analyst 2. To investigate the features of the tunnels during MD simulations, we analyzed the first 10 identified tunnel clusters ranked based on priority (calculated by averaging tunnel throughputs over all snapshots). Our calculations confirmed the persistence of the two dominant favorable tunnel clusters (1 and 2 in S5a Fig) with a slightly larger average and maximum bottleneck radius than similar tunnels initially calculated in the static X-ray structure. Both tunnel clusters were lined by predominantly hydrophobic residues as found in the case of static tunnels. This suggests that structural changes of LcpK30 do not largely influence the ranking and features of the putative pathways for substrate access. Interestingly, we also observed a new branched pathway connecting the heme co-factor in the protein interior with the surface through the tunnels surrounded by Glu148, Ile145, Asp226, Val229 and Thr230 (cluster 3 in S5a Fig). Due to the salt bridge between Arg147 and the neighboring Glu52 and Asp226, cluster 3 splits from its main route into two pathways. Because of its smaller average bottleneck radius, this cluster might be important for the transport of smaller products after the cleavage or O_2_ into and from the active site. We also noted that such a narrow tunnel could only be observed in the static structure after decreasing a minimum probe radius. The details about the features of the dynamic tunnel clusters are shown in S2 Table.

The analysis of potential protein cavities using the snapshots from MD simulations also confirmed the persistence of two hydrophobic pockets that could accommodate ligands in the active site near heme, both with slightly increased average volumes compared to the static structure. The larger pocket 3 directly interacts with heme and has an average volume of 182.08 Å^3^. A smaller pocket 12 has an average volume of 116.97 Å^3 and^ is buried deeper in the protein interior (S5b Fig). Interestingly, we observed a water network occupying the tunnels and both hydrophobic pockets above heme. These water molecules should be easily replaced by lipophilic ligands such as *cis*-1,4-polyisoprene.

### Molecular docking of the *cis*-1,4-polyisoprene substrate into LcpK30

Computational modelling of the interaction between flexible oligomeric ligands and their protein receptors is challenging. Dynamic modelling would be the best approach to account for the flexibility and obtain robust results [21,22]. Modelling of the interaction between a polyisoprene-type substrate and LcpK30 has been previously reported in an elegant study by Zhang and Liu, in which they carried out docking of a short oligomer with four repeating units into the binding site near the heme cofactor containing a bound dioxygen [16]. In their work, the authors primarily focused on the QM/MM calculations of the LcpK30 catalytic mechanism featuring the oxidative cleavage of the C=C bond. However, such a short substrate model does not provide the full information about the interactions occurring further away from the active site, for example within the tunnels and hydrophobic pockets, that could additionally influence its binding and the catalysis outcome. On the other hand, the natural substrate of LcpK30 is a long polymer consisting of more than 10,000 repeating isoprene units, which would be too complex for standard docking into protein [23]. The major oligomers produced in the reaction with LcpK30 typically range from C15 to C35 (5 to 7 repeating isoprene units). Therefore, we reasoned that a substrate model with 10 repeating isoprene units (C_50_H_82_) was sufficiently large to provide valuable insights into the interaction of the polymer substrate with the enzyme.

Our docking calculations were carried out in the GOLD software, using two different scoring functions, ChemPLP and ChemScore, which differ in the calculation of the fitness scores (see Methods). A higher fitness score indicates a higher probability that a ligand will bind to a protein in the given docking position [24]. The crystal structure of LcpK30 in its open conformation was used as the protein receptor, where a dioxygen molecule was manually coordinated to the Fe(II) in heme. The binding site included all atoms within 10 Å of radius from the central iron atom in heme, which also contained the residues reported to be part of the active site of LcpK30 (Arg164, Lys167, Thr168 and His198) [12]. In this initial standard docking approach, the substrate was fully flexible while the protein was kept rigid (we refer to this as rigid docking). However, by using the rigid docking approach, we found that even the highest ranked poses had a very low, or even negative fitness score, indicating that significant steric clashes occurred between protein and substrate. Therefore this docking approach was unsuitable to accurately describe the interaction between LcpK30 and the polyisoprene substrate (S6 Fig and S3 Table).

To improve docking quality, induced fit (flexible) docking was carried out, accounting for flexibility of the receptor by allowing a full torsional flexibility of 10 amino acid sidechains in the active site of LcpK30 (see Methods for details and Table 1 for docking results). Increasing the size of the defined binding site from 10 Å to 15 Å was tested but was not suitable; although higher fitness scores were obtained, the oligomeric substrate tended to dock preferentially at the surface of the protein, without interaction with the heme and the active site of LcpK30 (S4 Table and S7 Figure). The top 10 ranked structures obtained using induced fit docking with each of the ChemPLP and ChemScore protocols were significantly different from those obtained with rigid docking and showed higher fitness scores. To obtain further information on their orientation for catalysis, we analyzed the proximity of the C=C bond to the distal oxygen and the polymer conformation based on its span, its chain terminus position, and the position of the cleaved double bond (Table 1). In most docked poses, the distance between the C=C bond and the distal oxygen was below 5 Å, indicating that these structures could indeed represent pre-reactive states. Some exceptions were observed (ChemPLP docking pose rank 7; ChemScore docking poses 6 and 8), where the substrate was further away from the heme.

**Table 1.** Structural features of the best docking poses obtained from induced fit docking within GOLD. . Docking solutions are ranked based on the fitness score from highest to lowest.

| Rank | Fitness Score | C=C closest to Fe-O<sub>2</sub> complex<sup>a</sup> | Distance of C=C to the oxygen atom (Å)<sup>b</sup> |  | Ligand conformation |
| --- | --- | --- | --- | --- | --- |
|  |  |  | -H<sub>2</sub>C-C= | =C-CH<sub>3</sub> |  |
| ChemPLP docking protocol |  |  |  |  |  |
| 1 | 78.94 | 6 | 5 | 5.3 | Folded |
| 2 | 67.25 | 5 | 5.2 | 4 | Folded |
| 3 | 64.8 | 3 | 4.3 | 3.5 | Extended |
| 4 | 40.51 | 7 | 5.9 | 5.1 | Folded |
| 5 | 29.74 | 9 | 3.8 | 2.5 | Extended |
| 6 | 21.37 | 8 | 2.2 | 2.6 | Extended |
| 7 | 19.99 | 9 | 5.9 | 7 | Extended |
| 8 | -13.14 | 3 | 2.6 | 2.4 | Extended |
| 9 | -42.16 | 4 | 5 | 5.4 | Folded |
| 10 | -60.12 | 5 | 3.2 | 2.9 | Extended |
| ChemScore docking protocol |  |  |  |  |  |
| 1 | 32.65 | 6 | 5.8 | 4.5 | Folded |
| 2 | 29.29 | 8 | 3.3 | 3.8 | Extended |
| 3 | 18.16 | 4 | 4 | 3.1 | Extended |
| 4 | 13.57 | 3 | 3 | 2.9 | Extended |
| 5 | 12.1 | 3 | 2.9 | 2.6 | Extended |
| 6 | 5.8 | 7 | 8.3 | 9 | Extended |
| 7 | 5.44 | 8 | 3.4 | 3.1 | Extended |
| 8 | -2.2 | 1 | 7.8 | 7.2 | Extended |
| 9 | -3.92 | 3 | 3.7 | 3.3 | Folded |
| 10 | -6.98 | 8 | 4.1 | 3.9 | Extended |

<sup>a</sup> the first C=C unit number was counted from the end terminal with two methyl group R-C=C-(CH<sub>3</sub>)<sub>2</sub> – see scheme
<sup>b</sup> the distance was measured from each C atom in the double bond to the distal O atom of the Fe-O<sub>2</sub> complex

Overall, we found two prominent docked conformations of the oligomeric substrate, which we characterized as extended and folded. An extended conformation was defined as a structure in which one terminus of the oligomeric substrate model occupied the hydrophobic cavity within the protein interior near to heme (consistent with pocket 3 in our previous finding on the identification of tunnels and pockets), interacting with the residues above the heme at a distal site, whilst the other terminus was exposed to solvent at the surface of the protein. A folded conformation was defined where both termini were found towards the surface of the protein (see S8 and S9 Figs). In both these conformations, a C=C bond was found in the proximity of the distal oxygen, thus increasing the possibility of the reaction. The poses that were further away from the heme had an extended-like conformation. Interestingly, the higher docking scores using both ChemPLP and ChemScore functions were obtained with folded conformations. This probably occurred due to reduced steric clashes between atoms of neighboring residues in the protein structure, which increased the fitness scoring function for the folded conformation.

PLIP analysis of the substrate-LcpK30 contacts revealed mostly hydrophobic interactions. Three residues, Lys167, Leu171 and Ile396 were involved in interactions with all docked conformations, with Ala159 involved in interactions with most docked conformations obtained with both ChemPLP and ChemScore (see S5-S6 Tables and S10 Fig). Residue Lys167 has previously been reported to serve as a gating mechanism that opens and closes the entrance to the hydrophobic channel of the enzyme active site [10]. Our docking results infer that this residue might also play an important role in stabilizing protein-substrate complexes. Furthermore, only the extended docked conformations interacted with the Glu148 side chain, located further inside the hydrophobic channel above the heme and previously suggested to play a function in fine-tuning the active pocket to accommodate the substrate for reaction [16]. Earlier work on mutation of Glu148 to alanine, histidine, and glutamine revealed that the glutamate residue affected the specific activity of the enzyme but showed no significant differences in biochemical and biophysical properties, as well as no change in oligoisoprenoid product distribution [10]. Given this, we hypothesize that the extended substrate conformation is more representative of a catalytically relevant pose of the substrate within LcpK30. However, it cannot be excluded that the cleavage occurs with the substrate in the alternative, folded conformation, given the overall higher docking scores and flexibility of the system.

### Molecular dynamics simulation of LcpK30 in the presence of *cis*-1,4-polyisoprene

We further explored the influence of substrate binding on the conformation of LcpK30 by performing MD simulations of the enzyme in the presence of a docked oligomeric substrate. We carried out simulations for all top 10 ranked solutions obtained with ChemPLP and ChemScore fitness function (S11-S15 Figs). Compared to the case without the substrate, relatively larger RMSD values of the enzyme C-alpha atoms were observed in the presence of the substrate, sometimes even above 2 Å, especially in the case of the substrates in the extended conformation that bind deeper within the protein interior, such as docking solutions 10 and 3 from ChemPLP and ChemScore, respectively. From the RMSF calculations of the C-alpha atoms during the simulation of LcpK30 with the substrate bound, we observed overall very similar fluctuations as in the case of the enzyme without the substrate, which could indicate that these are characteristic for the protein in solution irrespective of whether substrate is present or not. However, when comparing the high RMSD docking solutions with the structure without the substrate, we characterized an increased fluctuation of protein residues 145-154 from a short helix interacting with the substrate bound in the hydrophobic pocket near the heme cofactor.

Moreover, larger fluctuations were also observed for the gateway loop residues 155-160 and the helix E residues 160-176, containing the flexible Lys167 (see S11a and S11b Figs for RMSD and RMSF plots of LcpK30 with docked substrates obtained with ChemPLP and ChemScore, respectively). Whilst the enzyme structure was not significantly affected by the substrates bound in the folded conformation, the majority of these newly appearing fluctuations of LcpK30 were caused by substrates in the extended conformation that bind deeper into the protein. To better visualize the atomic fluctuations, in S12 and S13 Figs of the SI we projected the RMSF values on the backbone of LcpK30. Overall, these results indicate that, for the substrate to bind in the hydrophobic pocket of the active site, subtle mutual structural rearrangements of the LcpK30 receptor, especially around the gateway, and of the polymeric chains are crucial for the formation of the stable complex.

Following the protein dynamics, we investigated the conformations of the substrate bound in the enzyme during MD simulations, to gain more insight into the stability of the complex initially obtained with docking. While we observed minor fluctuations of the polymeric chain that binds deeper into the protein and tightly interacts with the heme cofactor, no significant conformational changes of the substrate occurred during 100 ns simulations, compared to the initial structures obtained with docking. Solvent-exposed regions experienced more flexibility (see S14 and S15 Figs for superimposed MD snapshots starting from ChemPLP and ChemScore docking poses, respectively). This could indicate that the interactions with the enzyme position and stabilize the flexible substrate in a suitable conformation for the reaction with the heme-bound oxygen.

To investigate the substrate-enzyme interactions, we calculated close contacts occurring in MD simulations and extracted information about the specific residues that most frequently interact with different ligand conformations (see S7 and S8 Table in the SI for details). Based on the interaction patterns, the two main types of bound conformations observed with docking (extended and folded) were confirmed. Similar to the docking results, some substrates were found in an extended-like conformation, without interactions with the hydrophobic cavity. The simulation confirmed that the distinctive difference between the extended and folded conformations arises from the fact that ligands in the extended conformation can reach the buried hydrophobic protein cavity, which is not the case for folded structures. Both conformations readily interacted with heme and O_2_. The most frequent interactions and representative conformations of the substrate model bound in the active site of LcpK30 are shown in Figure 3.

**Fig 3.**
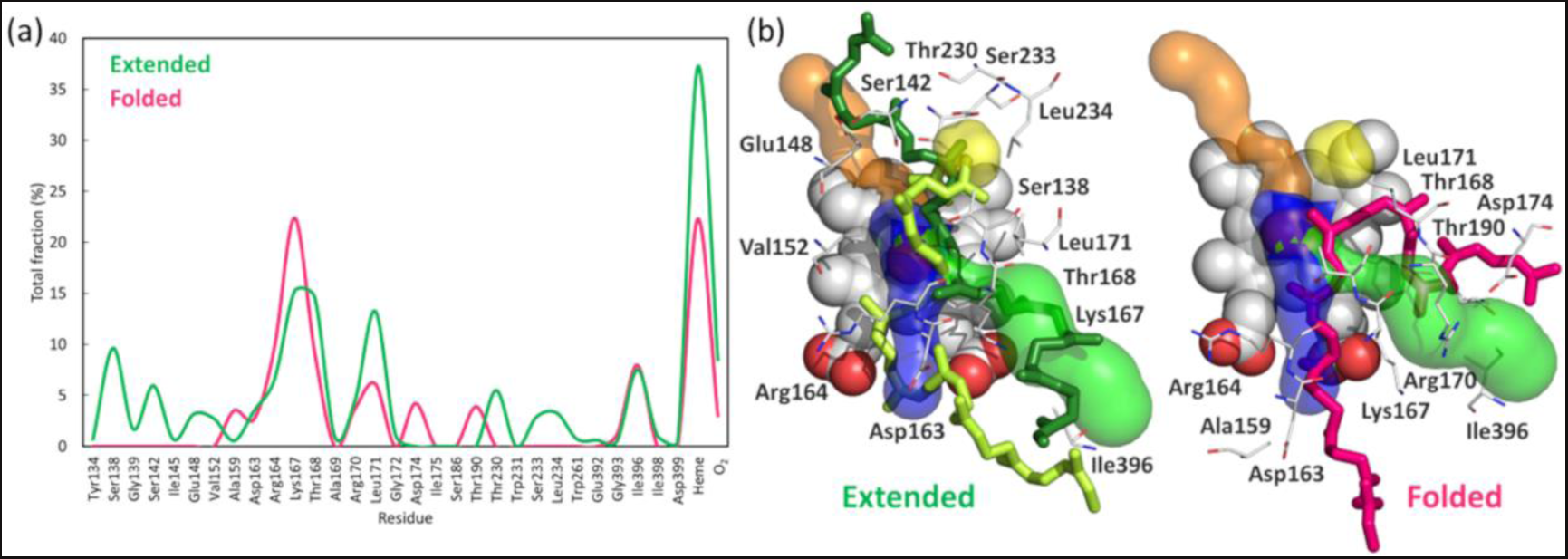
Residues that frequently interact with the substrate. (a) Close contacts between frequently occurring bound conformations of the substrate (extended and folded) and LcpK30, calculated from 100 ns MD simulations of 20 docking poses obtained with ChemPLP and ChemScore fitness function. The profile was constructed by averaging total fractions obtained for similar conformations. Conformations were clustered based on the contacts and the features from the visual inspection; (b) Representative snapshots extracted from MD simulations showing the main extended (ChemPLP pose 10 and ChemScore pose 5 shown as dark green and light green licorice, respectively) and folded (ChemScore pose 1 shown as magenta licorice) conformations and protein residues (shown as lines) that interact with the *cis*-1,4-polyisoprene substrate model bound to LcpK30 near heme (sphere representation). The three tunnels and the hydrophobic cavity are shown as blue, green, orange and yellow surface, respectively. The rest of the LcpK30 and hydrogen atoms are omitted for clarity.

Analyzing the frequent close contacts with the protein and the binding features of the extended and folded conformations of the *cis*-1,4-polyisoprene substrate bound to LcpK30 during MD simulations, we observed that both conformations readily interacted with the two previously found putative protein tunnels, as depicted in Fig 3. The main differences between the two binding modes arise from the fact that only the substrate in the extended conformation could reach the buried hydrophobic cavity, which allows it to favorably interact with the hydrophobic residues in the protein interior and positions a double bond near the heme-bound oxygen. Interestingly, we also found extended conformations that bound even deeper inside the protein, allowing the substrate termini to interact with a narrow tunnel 3. In such conformations, one part of the polymeric chain was bound in the active site while the other was exposed to the surface interacting with either tunnel 1 or 2. In contrast, the substrate in the folded conformation did not reach the hydrophobic pocket but rather coiled around Lys167 and the flexible C-terminus loop comprised of residues 392-397, with a central double bond positioned near the heme active site in an orientation favorable for catalysis.

To expand our investigation, we explored dominant energy factors that drive the *cis*-1,4-polyisoprene binding to LcpK30, by calculating van der Waals and electrostatic energy terms with linear interaction energy [25] on structures of the complex from MD simulations of all docking solutions from ChemPLP and ChemScore. Overall, we found that E_vdW_ dominates the total interaction energy with 94% contribution over E_ele_, which contributes only 6% as shown in S16 Fig of the SI. Furthermore, the binding energy was calculated with the MM/GBSA method [26] using snapshots from MD simulations and treating the solvation implicitly. Interestingly, the calculated binding energies show similar trends as observed for the total interaction energies. Whilst most docking solutions, irrespective of the conformation, have similar binding energy, structures 9 and 10 from the ChemPLP fitness score have significantly higher or lower binding energy, respectively. We further analyzed the relationship between interaction energies, and therefore binding affinities, and the total number of close contacts between the substrate and LcpK30 during MD simulations. We found that the binding affinity decreases when there are too few or too many close contacts. For example, the conformer with the most favorable binding affinity had 9 frequent contacts with the protein residues, while the conformer with the least favorable binding had only two contacts less. However, the conformation that achieved the least interactions with the protein (5 close contacts) had less favorable binding energy than the structure that had the most interactions (15 contacts). Moreover, we observed that the substrates in the extended conformation typically achieved 9 or more frequent contacts with the protein residues, while this number is less than 9 close contacts in the case of substrates that bind in the folded conformation (see S7 and S8 Tables in the SI for close contacts analysis on snapshots from ChemPLP and ChemScore, respectively). It is important to mention that the calculated binding energies do not correlate with the docking fitness scores, which highlights the importance of the flexibility of the system and the advantage of exploring different bound conformations of a ligand with molecular simulations.

The cleavage sites in the *cis*-1,4-polyisoprene substrate model were predicted by constructing the histograms of distances calculated between distal oxygen atom from heme-O_2_ cofactor and each of the ten C=C double bonds of the substrate during MD simulations. The near-attack-conformations of the substrate, in which the C=C bonds are frequently found around or below 4 Å from the O_2_, were considered suitable for the oxidative cleavage by LcpK30, as shown in S17 and S18 Figs for structures originating from ChemPLP and ChemScore fitness function, respectively. We used this information to construct the oxidative cleavage frequency profile by assigning the total number of potential cleavage events to a specific C=C bond based on their distance from the reactive O_2_. Fig 4 shows that the central C=C double bonds (numbered 5 and 6) of the substrate were more frequently found in the near vicinity of heme-O_2_ and therefore more prone to oxidative cleavage, which agrees well with the experimentally observed distribution of oligomers produced by LcpK30. It is important to mention that 4 out of the 6 folded conformations of the substrate had central C=C bonds in a favorable position for cleavage. On the other hand, the oxidative cleavage of the substrates in the extended conformation seemed to be more diverse and preferably occurred at the C=C bonds 3, 4 and 8 (see S17 and S18 Figs in the SI).

**Fig 4.**
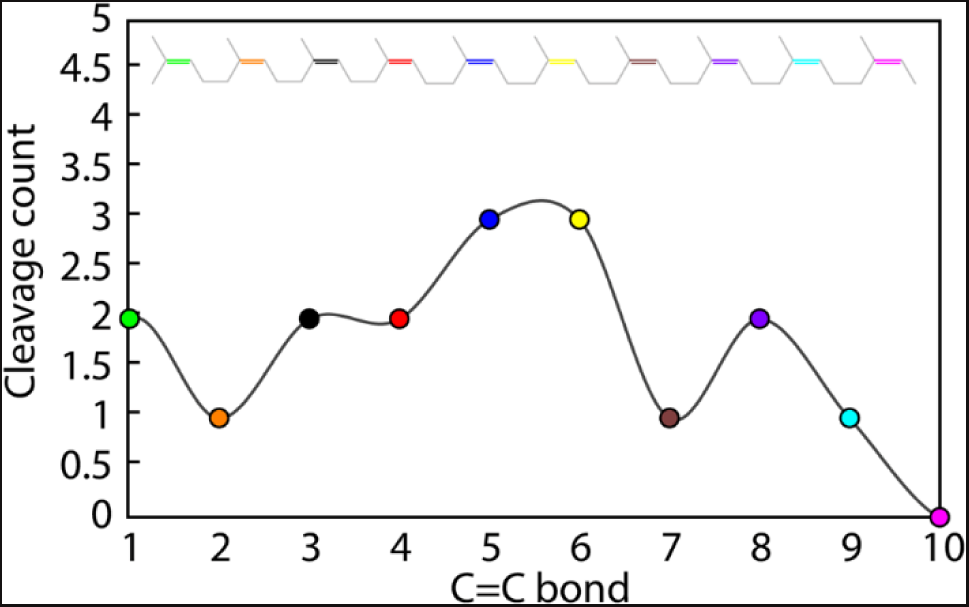
Cleavage frequency profile. Cleavage frequency profile constructed from distance histograms indicating a total count of each of the ten C=C double bonds (coloured differently) from the *cis*-1,4-polyisoprene substrate that are found in the near vicinity (below 4 Å) of the distal oxygen atom in the heme-bound O_2_ molecule. The profile is calculated from 100 ns MD simulations of LcpK30 in the presence of the substrate for all 20 docking poses obtained with ChemPLP and ChemScore fitness functions.

## Conclusion

Knowledge about pathways that lead from the surface to the active site is important to understand enzymatic catalysis. Frequently, the residues comprising these pathways actively transport substrates and products between the active site and bulk solvent, which is particularly important for enzymes with buried active sites [27–29]. In this work, docking and molecular dynamic simulations provided a deeper understanding of the interaction between the *cis*-1,4-polyisoprene and the enzyme latex clearing protein, including a detailed characterization of the tunnels and hydrophobic pockets in LcpK30. Using the CAVER tunnel identification and the automated fpocket tools within the static LcpK30 structure, as well as MD simulation analyses, we characterized potential entry and exit pathways, as well as hydrophobic pockets for the binding of oligomeric *cis*-1,4-polyisoprene chains within the enzyme. A range of residues interacting with the substrate were identified and comprised Ser138, Ser142, Glu148, Val152, Arg164, Lys167, Thr168, Leu171, Thr230, Ser233 and Leu234. We suggest that these residues represent suitable targets for mutagenesis aimed at improving enzymatic degradation rates and/or substrate and oligoisoprenoid product range of LcpK30.

Computational docking of a large substrate model with 10 repeating isoprene units (C_50_H_82_), using two different scoring functions within the GOLD program, revealed two main bound conformations: extended and folded. Residues Lys167, Leu171, Ile396 and Ala159 were involved in the protein-substrate interactions in most of the docking solutions, and we suggest they play an important role in stabilizing protein-substrate complexes. Only the extended substrate conformations interacted with residue Glu148, which was previously shown to affect enzyme activity. Therefore, we suggest that the extended conformation is more representative of a catalytically relevant substrate pose within the enzyme. Further analysis using MD simulations revealed additional close contacts between the substrate in different conformations and the enzyme. This analysis also confirmed that hydrophobic (van der Waals) interactions were mostly responsible for mutual subtle conformational changes of both LcpK30 and the polymeric substrate upon binding. Furthermore, an oxidative cleavage frequency profile was constructed, which agreed well with the experimentally observed distribution of oligomers produced by LcpK30.

Taken together, the tunnel and pocket predictions, docking results and MD simulations suggest that the substrate enters *via* a gateway comprised of flexible loop 155-160 that extends to the helix 160-176 and the C-terminus residues 392-397 which could also serve as the potential exit pathway for the oligoisoprenoid products (Fig 5). A hydrophobic pocket exists above the heme, where the substrate binds, and which is likely to determine the size distribution of the oligomers. An optimal number of hydrophobic interactions favors better substrate binding, which suggests the substrate needs to bind deeper into the protein cavity for the reaction to occur and may explain the oligomer sizes obtained during cleavage. The most favorable substrate binding occurs in an extended conformation, with one terminus within the hydrophobic pocket and the other terminus at the surface of the protein. A second binding mode with the substrate in a folded conformation also occurred, with a central double bond in proximity to the heme-bound oxygen, leading to a preference for oligomers with 5 to 6 C=C bonds, as shown by experimental data.

**Fig 5.**
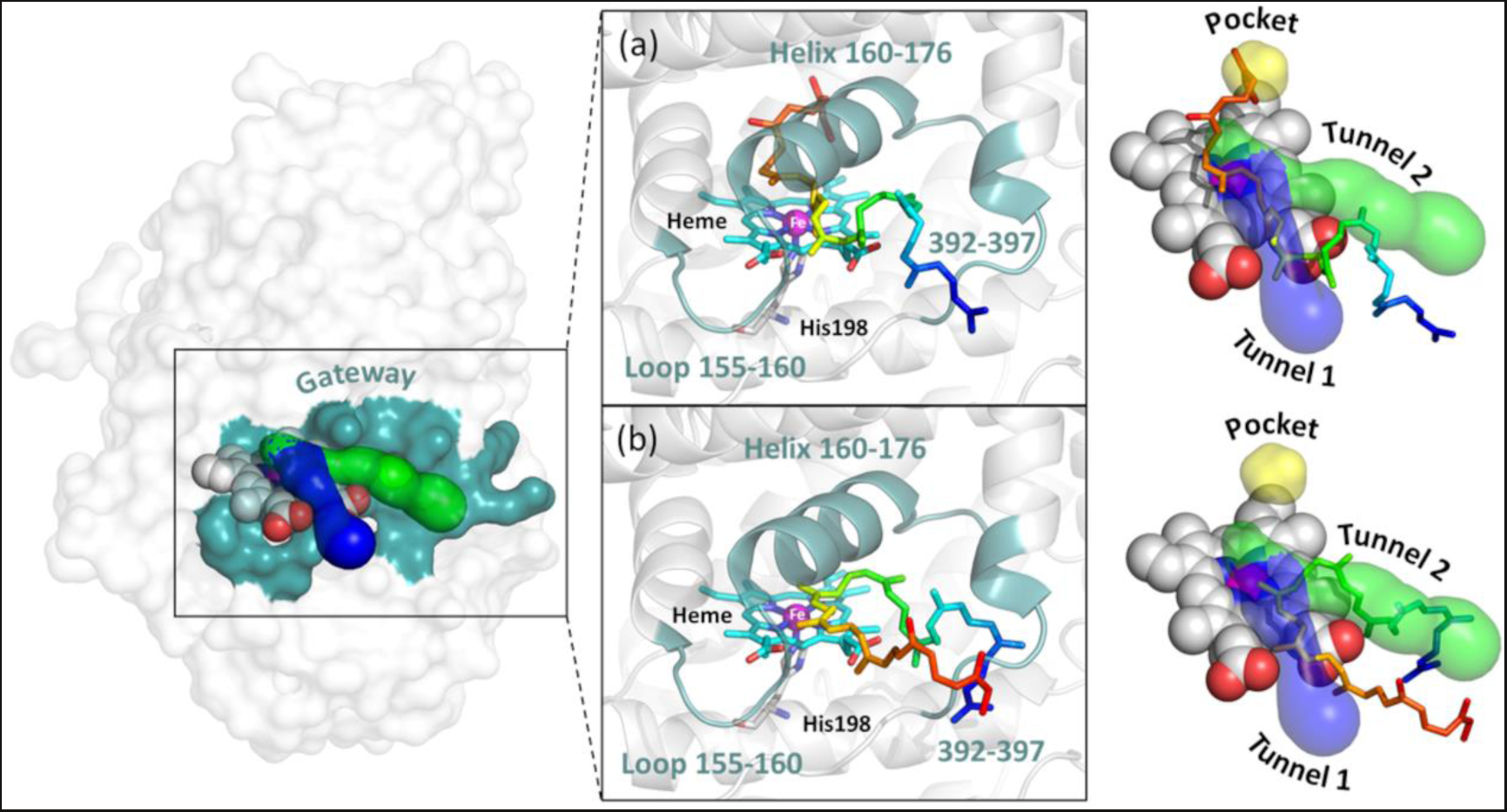
Proposed gateway for the substrate entry into LcpK30. The gateway residues were proposed based on the results from static and dynamic modelling of LcpK30 with *cis*-1,4-polyisoprene ligand. The representative snapshots of (a) extended and (b) folded conformation of *cis*-1,4-polyisoprene in complex with LcpK30 highlighting their interaction with the putative tunnels and the hydrophobic protein cavity. The O_2_ bound on heme was omitted for the sake of clarity.

The thorough computational investigation on the structure, interaction, and catalytic mechanism of LcpK30 discussed in this work will provide deeper insight into the understanding of enzymatic catalysis with other large, flexible and hydrophobic macromolecules.

## Computational methods

### Protein structure preparation

The high-resolution crystal structure of the LcpK30 enzyme in the open conformation was obtained from the RCSB Protein Data Bank (PDB ID: 5O1L) [10]. Although the LcpK30 is a biologically active dimer, only one monomer (chain A) was retained for all calculations and modelling because the interface between the two monomers is relatively far from the active site. The corresponding heme cofactor was also retained, while all other co-crystallized species including imidazole, 2,3-butanediol and 1,2-ethanediol molecule and waters were removed prior to all calculations.

### Protein tunnel calculations

Putative tunnels were calculated with the CAVER-Pymol plugin 3.0.3. [19]. The hydrogen atoms were added to the protein structure using Chimera 1.16, considering the H-bonds network [30]. The initial starting point for the tunnel search was specified by the central iron atom of the heme prosthetic group. The search was performed using a minimum probe radius of 0.9 Å. Default parameters for the maximum distance of 3 Å, desired radius of 5 Å, shell depth of 4 Å, and shell radius of 3 Å were used. The clustering of tunnels was performed by the average-link hierarchical Murtagh algorithm based on the calculated matrix of pairwise tunnel distances. The clustering threshold was set to a default value of 3.5. Each atom of the protein was approximated by 12 spheres. Calculations of tunnels on snapshots from MD simulations was carried out in CAVER Analyst 2 using default parameters [31].

### Protein pocket calculations

Simple pocket detection was performed with the open source protein pocket (cavity) detection algorithm based on Voronoi tessellation known as fpocket [32]. The detection of pockets on MD trajectories of LcpK30 was carried out with mdpocket considering protein and heme atoms only [33]. Default values of parameters to describe pockets were used. Namely, the minimum and the maximum radius that an alpha sphere might have in a binding pocket was 3.4 Å and 6.2 Å, respectively. The pairwise single linkage clustering method with the Euclidean distance measure was used for the clustering algorithm. Each pocket contained at least 15 alpha spheres and pockets volume were calculated using a Monte-Carlo algorithm.

### Protein-ligand docking

The GOLD program (version 2022.2.0) was used for the molecular docking protocols, with the default parameters as defined within the program [34]. The protonation states of the titratable residues were assigned using the H++ server [35]. To simulate the catalytically active state of the protein, the imidazole molecule coordinated on the iron of heme was replaced with O_2_ that was manually added. The 2D structure of the C50 *cis*-1,4-polyisoprene ligand model was constructed using ChemDraw 18.2 (Perkin Elmer, MA, USA) and the 3D structure with all hydrogen atoms included was obtained using Chem3D 18.2 (Perkin Elmer, MA, USA).

The ChemPLP and ChemScore fitness functions available in GOLD were used [24]. These scoring functions were selected because they both perform well for lipophilic binding sites and accurately model the steric complementarity between protein and ligand. All docking runs were performed with default parameters using both rigid and flexible docking. The binding site was defined as all protein residues whose atoms are within 10 or 15 Å from the central iron of heme. Ligand flexibility was enabled during all docking runs. To include flexibility of the active site, induced fit (flexible) docking was carried out, where the side chains of all residues that have been identified as important to the catalytic activity of LcpK30 (Glu148, Arg164, Lys167, Thr168) were specified as fully flexible [12]. Several other residues which were found in the pathway identified from the static tunnel calculations using CAVER (Ser142, Ile145, Val152, Leu171, Thr230 and Leu234) were also defined as flexible. These side chains were allowed to rotate freely during docking to vary over the range −180 to +180 rotatable torsion degree. The protein-ligand interaction was analyzed using Protein-Ligand Interaction Profiler (PLIP) program [36].

### Molecular dynamics simulations

The protonation states of the titratable residues at the physiological conditions were assigned using the H++ server [35]. The enzyme coordinates were taken from the crystal structure of the open (PDB ID: 5O1L) conformation of LcpK30, chain A.[10] The classical ff19SB [37] force field parameters were assigned to the standard residues, while the bonded model was used to describe the oxyheme cofactor and the coordinating His198. The cofactor was parametrized with the metal center parameter builder (MCPB) protocol [38]. To generate bonded force field parameters of the heme group with MCPB.py, we used a crystal structure of LcpK30 in the open state (PDB: 5O1L) [10] in which the central iron is coordinated by four equatorial nitrogen atoms from the protoporphyrin ring and one axial epsilon-nitrogen atom from the histidine sidechain of the enzyme. We manually included an oxygen molecule bound to the remaining iron site. We applied the B3LYP/6-31G(d) level of theory to carry out the geometry optimization for both the small and large models. Apart from the heme cofactor and O_2_ molecule, the small model had the sidechain of the coordinating histidine, while the large model also included the histidine backbone. The force constants were calculated for the optimized small model using the same method. Prior to the Merz-Kollman RESP partial charges calculation we performed minimization of the large model allowing the relaxation of hydrogen atoms and of O_2_ to the optimal positions. The +2 charge was defined for the iron atom, and the total charge of metal site was −2, while the metal site had a possibility to adopt multiple electronic structures. We tested different spin states and concluded that the bond parameters are closer to the experimental crystal structure of oxymyoglobin in the singlet spin state. The ESP charges, and consequently RESP charges, were similar both in the case of low and high spin systems. We employed the Seminario method to obtain force field parameters using the ff19SB/GAFF force field and ChgModB to perform the RESP charge fitting. All QM calculations were carried out with Gaussian 16 program [39].

To explore conformational dynamics of the enzyme and the interactions of the enzyme with the ligands, we simulated LcpK30 in the absence and in the presence of polyisoprene ligand using previously characterized docking poses of C50 *cis*-1,4-polyisoprene with 10 C=C double bonds. The GAFF force field was used to describe bonded parameters of the ligands while the partial charges were calculated using the AM1-BCC method [40,41]. We solvated the systems with OPC [35] water using the truncated octahedron box with each atom of the solute at least 10 Å away from the edge of water surrounding the solute. All systems were neutralized by adding Na+ counterions. The 12-6 set of parameters for monovalent atomic ions for the OPC water model were used [42]. All systems were minimized for a total of 10,000 steps, first 5,000 steps of the steepest descent method and the conjugate gradient for the remaining cycles. In the first relaxation step, the temperature of the system was gradually increased from the initial temperature of 0 to a target temperature of 300 K over 100 ps and kept constant for another 100 ps of the NVT simulation. We employed the Langevin thermostat with the collision frequency of 2 ps for the temperature control. In the second step of the relaxation, the system was propagated at 300 K over 100 ps of simulation time at a constant pressure (NPT), allowing the box to relax using the Berendsen barostat. The weak positional restraints (5 kcal mol^-1^ Å^-2^) were applied to the solute in the relaxation phase. This last step of relaxation included the removal of all restraints on the solute for 2 ns of simulation time at a constant pressure. Finally, for LcpK30 without the ligand, three independent production runs were carried out at 300 K and the constant pressure for at least 500 ns of simulation time. The production simulations of LcpK30 with the ligand were carried out for 100 ns each. All docking poses obtained from ChemPLP and ChemScore fitness functions were simulated. The Particle Mesh Ewald approach was used for treating long-range electrostatic interactions and the non-bonded cut-off was set to 10 Å. The SHAKE algorithm was used to constrain bonds involving hydrogen and a time step of 2 fs was used in all simulations. All MD simulations and the trajectory analysis were carried out in the AMBER20 program [43]. The visualization was performed in Pymol 2.5.4 [44] and VMD 1.9.3. [45].

## Acknowledgement

We would like to thank Prof. Derek Irvine for his contribution and useful discussion on project direction.

## Data Availability Statement

Input and output information from the computational analysis including Caver and fpocket setups; GOLD docking protocols; additional analysis from MD simulations and full MD inputs; and details on protein–ligand docking results are freely available on Figshare: https://doi.org/10.6084/m9.figshare.22941149.

## Author Contributions

Aziana Abu Hassan and Marko Hanževački conceived the analysis, collected data, performed the analysis and contributed to the writing. Anca Pordea and Marko Hanževački conceived and designed the analysis and wrote the paper. AP and MH are both corresponding authors.

## Supporting information

S1 Fig. Crystal structure analysis.

**S2 Fig. Tunnel profiles for the dominant tunnels. S3 Fig. The RMSD and RMSF values.**

**S4 Fig. (a) Experimental B-factor from the X-ray structure and (b) C-alpha RMSF (Å) values from MD simulations plotted on the backbone of LcpK30.**

**S5 Fig. Dominant superclusters of tunnels and pockets.**

**S6 Fig. Superimposed conformations results from rigid docking of cis- 1,4-polyisoprene C50H82 within LcpK30.**

**S7 Fig. Superimposed docking conformations results of C50H82 within a defined radius of (a) 10 Å and (b) 15 Å from the central iron atom in heme using ChemPLP flexible docking.**

S8 Fig. Docking conformations of cis-1,4-polyisoprene (C50H82) near heme, obtained with the ChemPLP fitness function.

**S9 Fig. Docking conformations of cis-1,4-polyisoprene (C50H82) near heme, obtained with the ChemScore fitness function.**

**S10 Fig. Protein-ligand interactions identified by PLIP analysis.**

**S11 Fig. C-alpha RMSD and RMSF calculated during 100 ns MD simulations of LcpK30 with cis-1,4-polyisoprene substrate model bound, starting from 10 docking poses obtained with (a) ChemPLP and (b) ChemScore.**

**S12 Fig. C-alpha RMSF calculated during 100 ns MD simulations of LcpK30 with cis-1,4-polyisoprene bound, starting from 10 docking poses obtained with ChemPLP fitness function.**

**S13 Fig. C-alpha RMSF calculated during 100 ns MD simulations of LcpK30 with cis-1,4-polyisoprene bound, starting from 10 docking poses obtained with ChemScore fitness function.**

**S14 Fig. Superimposed snapshots from 100 ns MD simulations of cis-1,4- polyisoprene dokcing on LcpK30 obtained with ChemPLP fitness function.**

**S15 Fig. Superimposed snapshots from 100 ns MD simulations of cis-1,4- polyisoprene dokcing on LcpK30 obtained with ChemScore fitness function.**

**S16 Fig. Linear interaction energy analysis between the ligand and LcpK30.**

**S17 Fig. Probability distribution of distance between distal oxygen of O2 and each C=C double bond. calculated during 100 ns MD simulations of LcpK30 with cis-1,4-polyisoprene bound, starting from 10 docking poses obtained with ChemPLP fitness score.**

**S18 Fig. Probability distribution of distance between distal oxygen of O2 and each C=C double bond, calculated during 100 ns MD simulations of LcpK30 with cis-1,4-polyisoprene bound, starting from 10 docking poses obtained with ChemScore fitness score.**

S1 **Table. The features of dominant tunnels calculated using the LcpK30 crystal structure in the open state (PDB 5O1L).**

S2 **Table. The features of the dominant tunnels calculated on 1500 snapshots from MD simulations of LcpK30.**

S3 **Table. Docking solutions obtained from rigid docking within GOLD software.**

S4 **Table. Impact of the size of the defined protein binding site on the docking results.**

S5 **Table. List of residues interacting with the docked cis-1,4-polyisoprene. extracted from PLIP analysis of 10 docking poses obtained using ChemPLP.**

S6 **Table. List of residues interacting with the docked cis-1,4-polyisoprene, extracted from PLIP analysis of 10 docking poses obtained using ChemScore.**

S7 **Table. Frequency of contacts between the substrate and LcpK30. calculated during 100 ns MD simulations of 10 docking poses obtained from ChemPLP fitness function.**

S8 **Table. Frequency of contacts between the substrate and LcpK30 calculated during 100 ns MD simulations of 10 docking poses obtained from ChemScore fitness function.**

